# PD-L1 expression is mediated by microRNA processing, Wnt/β-catenin signaling, and chemotherapy in Wilms tumor

**DOI:** 10.1101/2024.11.29.626084

**Authors:** Kavita Desai, Patricia D.B. Tiburcio, Austin Warne, Arash Nabbi, Serena Zhou, Sean D. Reiff, Matthew E. Campbell, Kenneth S. Chen

## Abstract

Inhibition of immune checkpoint proteins is effective in adult cancers but has shown limited efficacy in pediatric cancers. While factors regulating expression of immune checkpoint proteins such as PD-L1 are well-documented in adult cancers, their regulation is poorly understood in pediatric cancers. Here, we show that PD-L1 is upregulated in distinct subsets of Wilms tumor, the most common pediatric kidney cancer. Specifically, chemotherapy-exposed Wilms tumor specimens exhibited higher levels of PD-L1 expression, and common chemotherapeutics upregulated PD-L1 in childhood cancer cell lines *in vitro*. Furthermore, mutations in *CTNNB1* and *DROSHA*, the two most commonly mutated genes in Wilms tumor, correlated with higher PD-L1. Activation of Wnt/β-catenin signaling and knockdown of *DROSHA* or *DICER1* both increase PD-L1 *in vitro*. Lastly, in adult cancers, *DICER1* alterations are associated with immune gene expression signatures and improved survival in response to immune checkpoint inhibitors. Together, our results identify clinical and biological properties regulating PD-L1 in Wilms tumor that may inform precision therapy approaches in pediatric immuno-oncology.

## INTRODUCTION

Wilms tumor is the most common pediatric kidney cancer and accounts for 6% of childhood cancers^1^. About 90% of patients are cured with current treatment regimens, including surgery, chemotherapy, and radiation. However, high-risk features such as advanced stage, anaplastic histology, and chemorefractory disease continue to portend poor outcome, with survival around 50%^2^.

We previously showed that the most common recurrent Wilms tumor mutations fall into four classes, affecting microRNA processing (*DROSHA*, *DICER1*, *DGCR8*), kidney development (*WT1*, *CTNNB1*, *SIX1/2*), chromatin remodeling (*CREBBP*, *REST*), and MYCN (*MYCN*, *MAX*)^3^. Wilms tumors are diagnosed based on their characteristic “triphasic” histological pattern (blastema, epithelia, and stroma), which recapitulates the three types of cells seen in the developing embryonic kidney. Mutations in kidney development genes are thought to arrest cells in this embryonic state. Similarly, impaired production of microRNAs prevents the suppression of microRNA target genes, leading to a similar developmental arrest.

Immune checkpoint inhibitors (ICIs) have revolutionized adult cancer therapy, but it has been challenging to apply these advances to pediatrics, as limited responses to ICI monotherapy have been seen in children with solid tumors^4–6^. ICIs work by blocking signals that cancer cells use to evade the anti-tumor immune response, such as the tumor antigen programmed death ligand 1 (PD-L1), and PD-1, its cognate ligand on T cells^7–10^. Lack of response to ICIs has been attributed to the lower tumor mutational burden (TMB) seen in pediatric cancers (including Wilms tumor)^11,12^. Along these lines, while adult cancers with significant lymphocytic infiltrate are more likely to respond to ICIs, most pediatric cancers are “immune cold” tumors, with little inflammatory infiltrate^13^. In many adult cancers, chemotherapy can sensitize tumors to ICI or enhance their effect, even in cases with low PD-L1 expression or low TMB^7,8,14–18^. It is unknown whether chemotherapy can serve the same role in pediatric cancers. Furthermore, while genome-wide mutational burden clearly drives tumor immunogenicity, ICI response may also be dictated by individual mutations in genes that regulate chromatin remodeling (*ARID1A*, *PBRM1*, *SMARCB1*), RNA processing (*ADAR1*), or immune response (*B2M*, *JAK1/2*)^5,7,8,19–28^.

Here we show that certain subsets of Wilms tumor are also associated with PD-L1 upregulation. The most common Wilms tumor mutations, in *CTNNB1* or *DROSHA*, were associated with higher PD-L1. Suppression of microRNA processing in a Wilms tumor cell line also led to PD-L1 upregulation *in vitro*, and adult cancers with impaired microRNA processing exhibited immune expression signatures. Furthermore, chemotherapy-treated Wilms tumors exhibited higher levels of PD-L1, and chemotherapy also induced PD-L1 in multiple cell lines across different childhood cancers. In summary, we identify clinical and biological features of Wilms tumor that drive PD-L1 expression.

## RESULTS

### A subset of Wilms tumors marked by immune signaling

To understand how clinical and molecular features affect protein levels and post-translational modifications in Wilms tumor, we used reverse-phase protein arrays (RPPA) to analyze a set of 48 Wilms tumor samples that had previously undergone genomic analysis^29^ (**Suppl. Table S1**). RPPA is a targeted proteomics platform that quantifies hundreds of proteins and post-translational modifications (PTMs) in parallel^30^. By unsupervised clustering of normalized RPPA results, we found that Wilms tumors formed three distinct clusters (**Fig. 1A, Suppl. Table S2**). Clusters 1, comprising 13 tumors, was marked by high expression of immune signaling markers and immune regulatory proteins, such as phospho-NF-κB, phospho-Stat3, IL-6, PD-1, PD-L1, B7-H4. Cluster 2 had moderate expression of these immune markers, but higher levels of other immune markers (CD4 and STING), as well as phosphorylated S6, a marker of mTORC1 signaling. Cluster 3, comprised of 24 tumors, was marked by low expression of immune markers and higher expression of cell cycle regulatory proteins and DNA damage repair proteins, such as cyclin B1, MSH6, PARP, MSH2, and 53BP1, reflecting high levels of proliferation and DNA synthesis.

**Figure 1.**
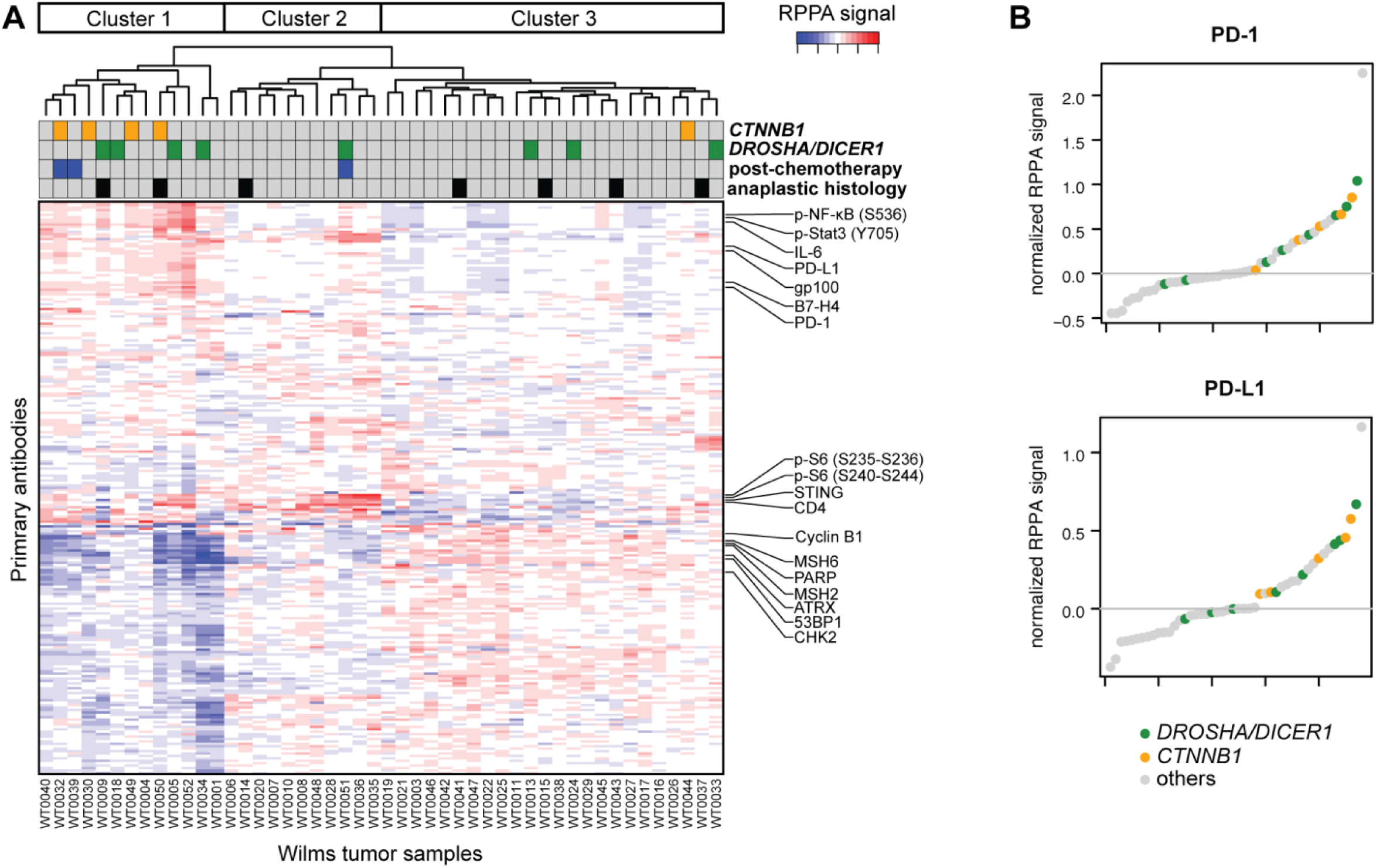
Protein array reveals a subset of Wilms tumors marked by immune signaling. (A) Unsupervised clustering of Wilms tumor clinical, genomic, and RPPA data reveals three distinct clusters with differential protein expression. (B) PD-1 and PD-L1 RPPA signal, highlighting tumors with mutations in *DROSHA/DICER1* or *CTNNB1*.

We examined clinical features that correlated with these clusters (**Fig. 1A**). Anaplastic histology is associated with particularly poor outcomes, and a previous report correlated anaplastic histology with PD-L1 overexpression by immunohistochemistry^31,32^. However, anaplastic histology did not correlate with PD-L1 upregulation or Clusters 1 and 2 in our dataset (**Suppl. Fig. S1A**). Very few Wilms tumor patients are treated with neoadjuvant chemotherapy in North America^1^, and only 3 samples in our cohort came from chemotherapy-exposed patients. Notably, all three chemotherapy-treated tumors were in Clusters 1 and 2 and exhibited higher levels of immune markers (**Fig. 1A, Suppl. Fig. S1B**).

We then examined molecular factors. Specifically, we interrogated the two most commonly mutated genes in Wilms tumor, *DROSHA* and *CTNNB1*^33^. Mutations in microRNA processing genes and *CTNNB1* were both more prevalent in Cluster 1, though only *CTNNB1* mutations reached statistical significance (**Fig. 1B**). Specifically, mutations in *DROSHA*/*DICER1* comprised 4 of 13 tumors in Cluster 1 and 4 of 35 tumors in the other two clusters (31% vs. 11%, p=0.25), while *CTNNB1* mutations were seen in 4 of 13 tumors in Cluster 1 and 1 of 35 tumors in the other two clusters (31% vs. 3%, p=0.02). Mutations in *CTNNB1* or *DROSHA/DICER1* were associated with higher levels of several immune markers (**Suppl. Fig. S2A-S2B**). Prior reports had suggested that copy number changes at the human leukocyte antigen (HLA) locus could be associated with cancer immune evasion^34,35^; however, we did not find copy number changes at this locus across our tumor cohort (**Suppl. Fig. S3A-S3B**).

We thus examined whether mutations in microRNA processing genes or in *CTNNB1* were associated with immune transcriptomic signatures in the published Therapeutically Applicable Research to Generate Effective Treatments (TARGET) cohort of high-risk Wilms tumor patients treated in the U.S.^33,36^. We used two different algorithms designed to detect immune infiltration in bulk RNA-seq data from primary tumor samples: “Estimation of STromal and Immune cells in MAlignant Tumor tissues using Expression data” (ESTIMATE)^37^ and “Quantification of the Tumor Immune contexture from human RNA-seq” (quanTIseq)^38^. Mutations in microRNA processing genes, but not mutations in *CTNNB1*, were associated with higher ESTIMATE immune scores (**Suppl. Fig. S4A-S4B**). We also used quanTIseq to identify signatures of individual immune cell types across these tumors. Mutations in *CTNNB1* were associated with a significantly increased population of dendritic cells, but we did not find increases in immune cell types among tumors with microRNA processing gene mutations (**Suppl. Fig. S4C-S4D**).

### Chemotherapy upregulates PD-L1 expression in Wilms tumor and other childhood cancers

We next examined how chemotherapy treatment affects tumor-immune interactions in an independent dataset. Specifically, we re-analyzed a published RNA-seq dataset^39^ that included Wilms tumors from both the U.S. (chemotherapy-naïve samples, n=120) and Europe (samples taken after neoadjuvant chemotherapy, n=17). Using Gene Set Enrichment Analysis (GSEA)^40^, we dissected how chemotherapy treatment affects gene expression pathways. The most significantly enriched “reactome” gene sets in chemotherapy-treated tumors were all immune-related, including “Immunoregulatory interactions between a lymphoid and a non-lymphoid cell” and “PD-1 signaling” (**Fig. 2A**, **Suppl. Fig. S5A**). Similarly, the most significantly enriched “hallmark” gene sets were immune-related, including “inflammatory response” and “TNF-ɑ signaling via NF--B” (**Suppl. Fig. S5B-S5C**). At the individual gene level, both *PDCD1* and *CD274* (PD-1 and PD-L1, respectively) were significantly overexpressed in chemotherapy-treated tumors compared to the chemotherapy-naïve cohort (**Fig. 2B**).

**Figure 2.**
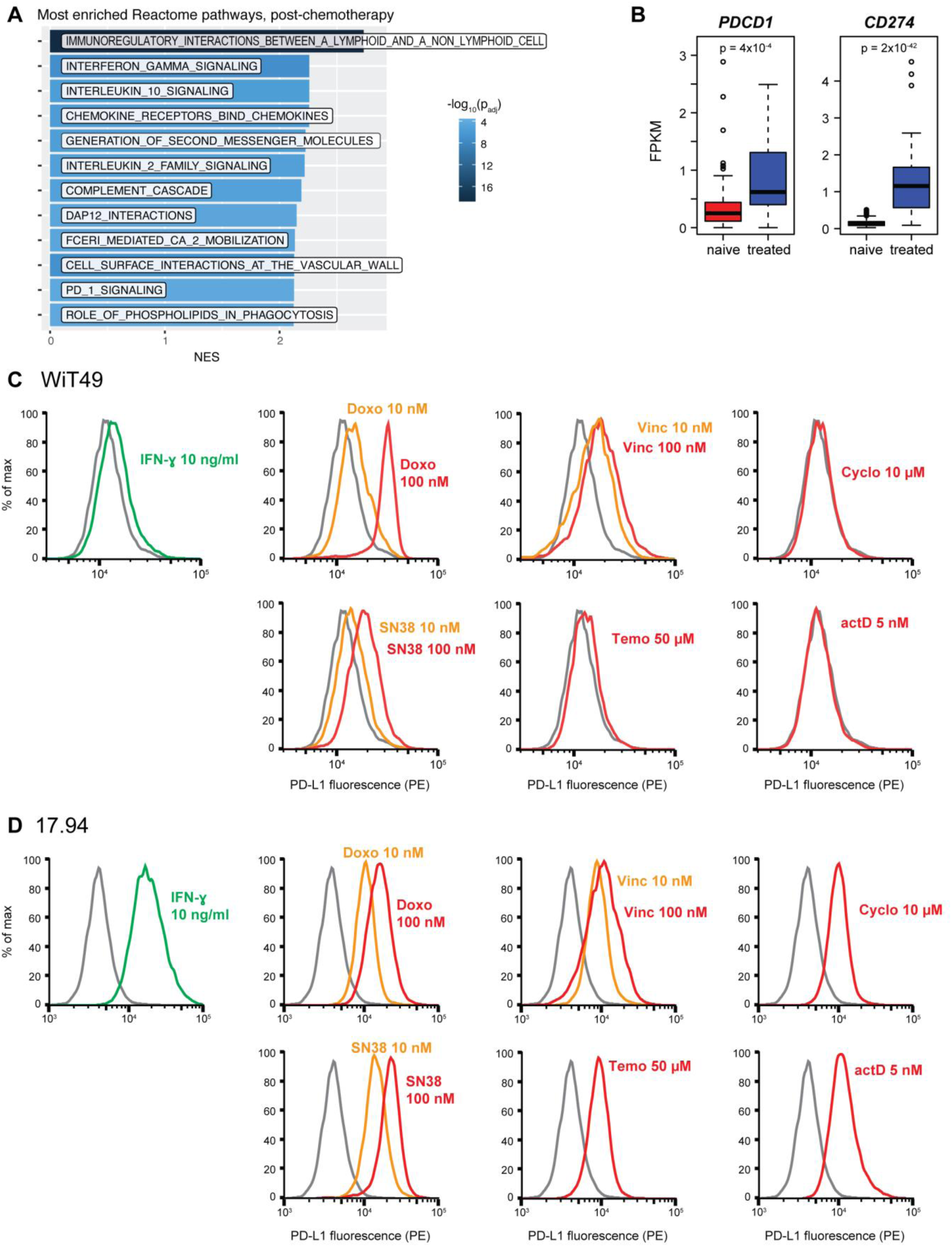
Chemotherapy induces PD-L1 in Wilms tumor. (A) Enrichment of Reactome gene sets in chemotherapy-treated vs. chemotherapy-naïve Wilms tumor specimens. NES, normalized enrichment score. (B) Expression of *PDCD1* and *CD274* in chemotherapy-treated vs. chemotherapy-naïve Wilms tumor specimens. FPKM, fragments per kilobase per million reads. (C) Flow cytometry data showing the effect of various chemotherapeutic agents on the expression of PD-L1 in Wilms tumor cell lines, WiT49 (C) and 17.94 (D). DMSO (gray) was used as negative control and IFN-γ (10 ng/mL) was used as positive control. IFN-γ, interferon gamma; Doxo, doxorubicin; Vinc, vincristine; Cyclo, cyclophosphamide; Temo, temozolomide; actD, actinomycin D.

We sought to validate this correlation *in vitro* by testing whether commonly used chemotherapy induces PD-L1 upregulation in the Wilms tumor cell lines WiT49 and 17.94. Specifically, we measured surface PD-L1 expression by flow cytometry after treatment with sublethal doses of the most common chemotherapy drugs used in Wilms tumor: doxorubicin, vincristine, cyclophosphamide, SN-38 (the active metabolite of irinotecan), temozolomide, and actinomycin D (**Fig. 2C**). In WiT49, doxorubicin, vincristine, and SN-38 induced dose-dependent increases in PD-L1, while actinomycin D, cyclophosphamide, and temozolomide, appeared to have little effect. In 17.94, positive shifts in PD-L1 expression were seen with all six drugs tested (**Fig 2D**). Overall, we found that the most common neoadjuvant chemotherapy regimens used in Wilms tumor induce PD-L1 surface expression.

Next, we examined whether these chemotherapy drugs also induce PD-L1 expression in cell lines derived from other embryonic cancers. We tested two rhabdomyosarcoma cell lines, JR-1 and RD, and two neuroblastoma cell lines, Kelly and SHEP. We treated these four cell lines with the same six chemotherapeutic agents as described above, and we found similar results. Across these non-Wilms tumor cell lines, we found that doxorubicin, vincristine, and SN-38 again induced PD-L1 overexpression (**Fig. 3A-D**). Actinomycin D, cyclophosphamide, and temozolomide had no appreciable effect on PD-L1 expression.

**Figure 3.**
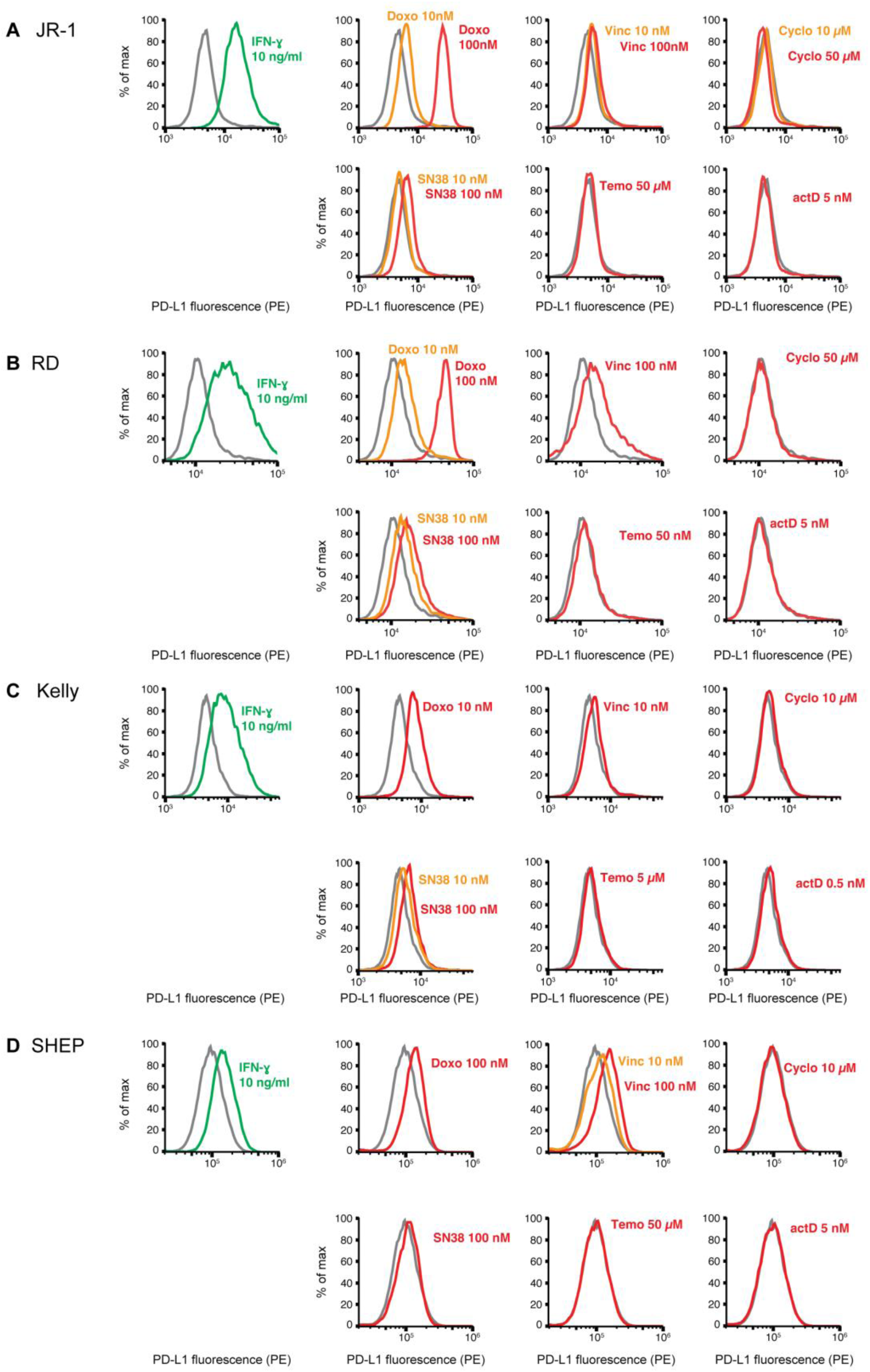
Chemotherapy induces PD-L1 in other childhood cancers. (A-D) Flow cytometry data showing the effect of various chemotherapeutic agents on the expression of PD-L1 in JR-1 (A), RD (B), Kelly (C) and SHEP (D). DMSO (gray) used as negative control and IFN-γ (10 ng/mL) was used as positive control. IFN-γ, interferon gamma; Doxo, doxorubicin; Vinc, vincristine; Cyclo, cyclophosphamide; Temo, temozolomide; actD, actinomycin D.

### Wnt/β-Catenin signaling activation upregulates PD-L1 temporarily in Wilms tumor cells

Because the results of our array data suggested that *CTNNB1* activating mutations were associated with increased PD-L1 and immune signatures, we examined whether Wnt/β-catenin activity upregulates PD-L1 expression in the Wilms tumor cell lines WiT49 and 17.94. While some studies have shown that β-catenin can directly activate PD-L1 transcription^43,44^, others suggest that high β-catenin activity in tumors is associated with decreased PD-L1 expression and reduced immune activation^45,46^. Thus, we treated our Wilms tumor cell lines with the small molecule CHIR-99021, which activates Wnt signaling by inhibiting glycogen synthase kinase 3 (GSK-3), the kinase that phosphorylates β-catenin and triggers its destruction^47^. Within three hours of CHIR-99021 treatment, we detected the accumulation of β-catenin and PD-L1 in both cell lines (**Suppl. Fig. S6A-S6B**). However, while both active β-catenin and total β-catenin continued to accumulate after 24 hours of treatment, PD-L1 appeared to decline at this time point. This suggests that Wnt/β-catenin signaling initially induces PD-L1 expression, but its expression wanes at later time points. This may partially explain the discrepant reports of the effect of Wnt/β-catenin signaling on PD-L1 expression in the literature. Nevertheless, our results support the assertion that Wnt/β-catenin signaling can, at least initially, upregulate PD-L1.

### DROSHA and DICER1 regulate PD-L1 in vitro

We next examined whether *DROSHA* and *DICER1* regulate PD-L1 in Wilms tumor cells. We previously used *DROSHA* silencing to model the impaired microRNA expression produced by dominant-negative *DROSHA* mutations seen in Wilms tumor^29,48,49^. To achieve stable knockdown in a microRNA-independent manner, we used CRISPR interference (CRISPRi), which uses single-guide RNAs (sgRNAs) to block transcription initiation at targeted regions^50,51^. In RNA-seq of *DROSHA*-silenced WiT49, we again observed upregulation of immune-related gene sets, including “Inflammatory response” and “TNF-ɑ signaling via NF-κB” (**Fig. 4A**). Next, we measured the activity of signaling pathways discovered from our Wilms tumor RPPA and RNA-seq analyses. We found that cells with either *DROSHA* or *DICER1* knockdown exhibit higher PD-L1 by Western blot, flow cytometry, and immunofluorescence (**Fig. 4B-4E**, **Suppl. Fig. S7A).**

**Figure 4.**
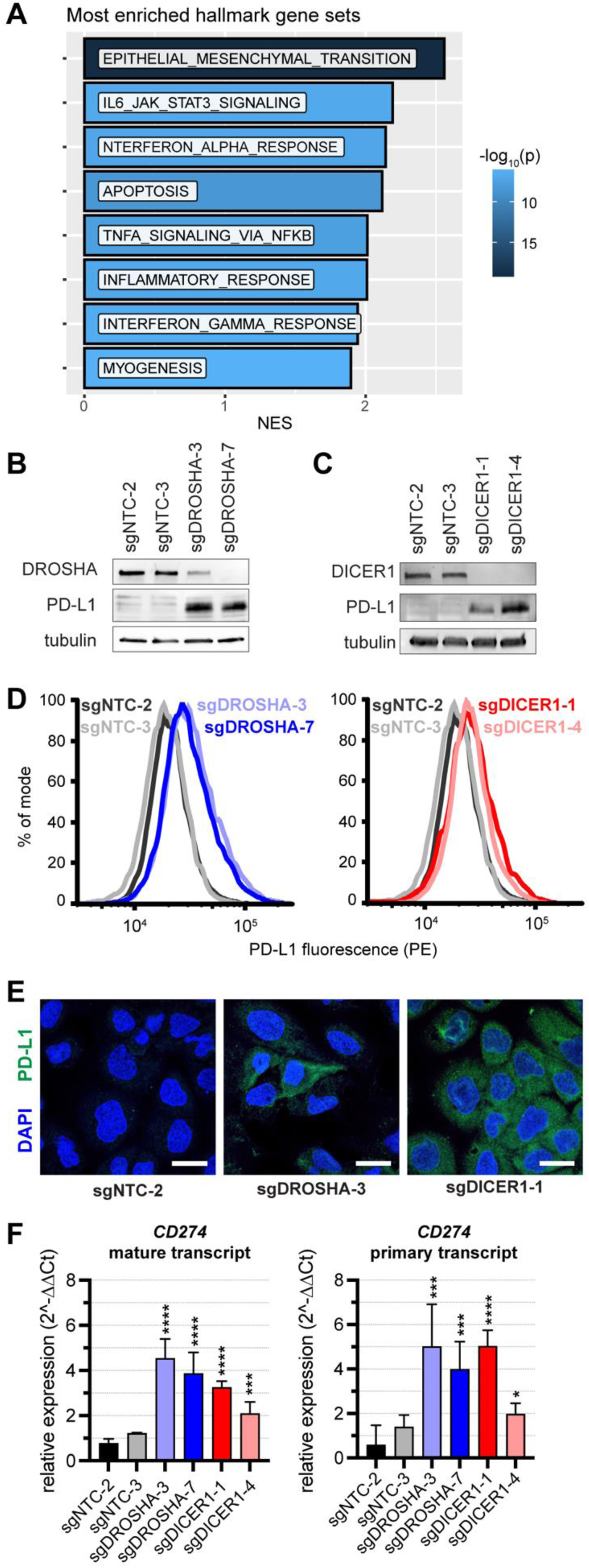
*DROSHA* and *DICER1* regulate PD-L1 in WiT49. (A) Most enriched hallmark gene sets in WiT49 with CRISPRi sgRNA against *DROSHA* (sgDROSHA) versus non-targeting control (sgNTC) cells. NES, normalized enrichment score. (B-C) PD-L1 Western blot of WiT49 with CRISPR interference knockdown of *DROSHA* (B) or *DICER1* (C), vs. non-targeting controls (NTC). (D) PD-L1 flow cytometry in WiT49 with knockdown of *DROSHA* or *DICER1* compared to non-targeting controls (NTC). (E) Representative images of PD-L1 immunocytochemistry of WiT49 *DROSHA* and *DICER1* knockdowns versus NTC (Scale bar = 25µm). See also **Suppl. Fig. S7**. (F) Normalized mature and primary *CD274* transcript levels by qPCR (****p<0.0001, ***p<0.001, *p<0.05, by unpaired two-tailed Student’s t-test versus NTC).

We explored whether PD-L1 accumulation in these cells was regulated at the transcriptional or post-transcriptional level. Loss of microRNAs leads to upregulation of microRNA target genes, which may lead to indirect upregulation of other genes. Because microRNAs repress their target genes post-transcriptionally, upregulation of direct microRNA targets is usually associated with an increase in a spliced, mature transcript without an increase in the unspliced, primary transcript. An increase in *CD274* (PD-L1) transcription, such as in response to interferon, results in an increase in both the mature and primary *CD274* transcripts (**Suppl. Fig. S7B**). We found that *DROSHA-* and *DICER1*-knockdown cells exhibited upregulation of both the pre-mRNA and mature mRNA, suggesting some transcriptional regulation (**Fig. 4F**).

Next, we examined whether the accumulation of abnormally processed microRNA precursors could account for the PD-L1 upregulation we observed. It had previously been suggested that double stranded RNA (dsRNA) activates the innate immune system and upregulates PD-L1 in response to knockdown of *DROSHA* or *DICER1*^52–55^. Thus, we examined whether we could detect dsRNA by immunofluorescence in *DROSHA/DICER1*-silenced cells. While NTC cells were negative for dsRNA, we found isolated dsRNA punctae by immunocytochemistry in some *DROSHA-* and *DICER1-*silenced cells (**Suppl. Fig. S7C**). *DROSHA* and *DICER1* have also been shown to regulate immunogenicity by regulating *Alu* RNA and other transposable elements, which can trigger innate immune signaling mechanisms^56–59^. We thus measured *Alu* RNA in *DROSHA/DICER1*-knockdown cells and found *Alu* RNA to be elevated in three of the four conditions (**Suppl. Fig. S7D**). In sum, we find that loss of microRNA processing induces PD-L1 upregulation through an indirect effect in Wilms tumor cells *in vitro*.

### Loss of microRNA processing corresponds with ICI response in adult cancers

Lastly, we investigated whether *DICER1* mutations correlate with immune gene signatures in adult cancers using publicly available RNA-seq from The Cancer Genome Atlas (TCGA). We first examined endometrial cancer^60^, the adult cancer most associated with recurrent oncogenic *DICER1* mutations^61^. As in Wilms tumor, immune gene sets were among the most enriched “reactome” gene sets, including “immunoregulatory interactions between a lymphoid and a non-lymphoid cell”, “PD-1 signaling”, and “CD28 co-stimulation” (**Fig. 5A**, **Suppl. Fig. S8A**). Similarly, two of the most enriched “hallmark” gene sets were “allograft rejection” and “interferon gamma response” (**Fig. 5B**, **Suppl. Fig. S8B**). Expression of *CD8A*, *PDCD1*, and *LAG3* were all significantly higher in *DICER1*-mutant endometrial cancers (**Fig. 5C**). In other words, *DICER1* mutations are also associated with immune signatures in adult endometrial cancer.

**Figure 5.**
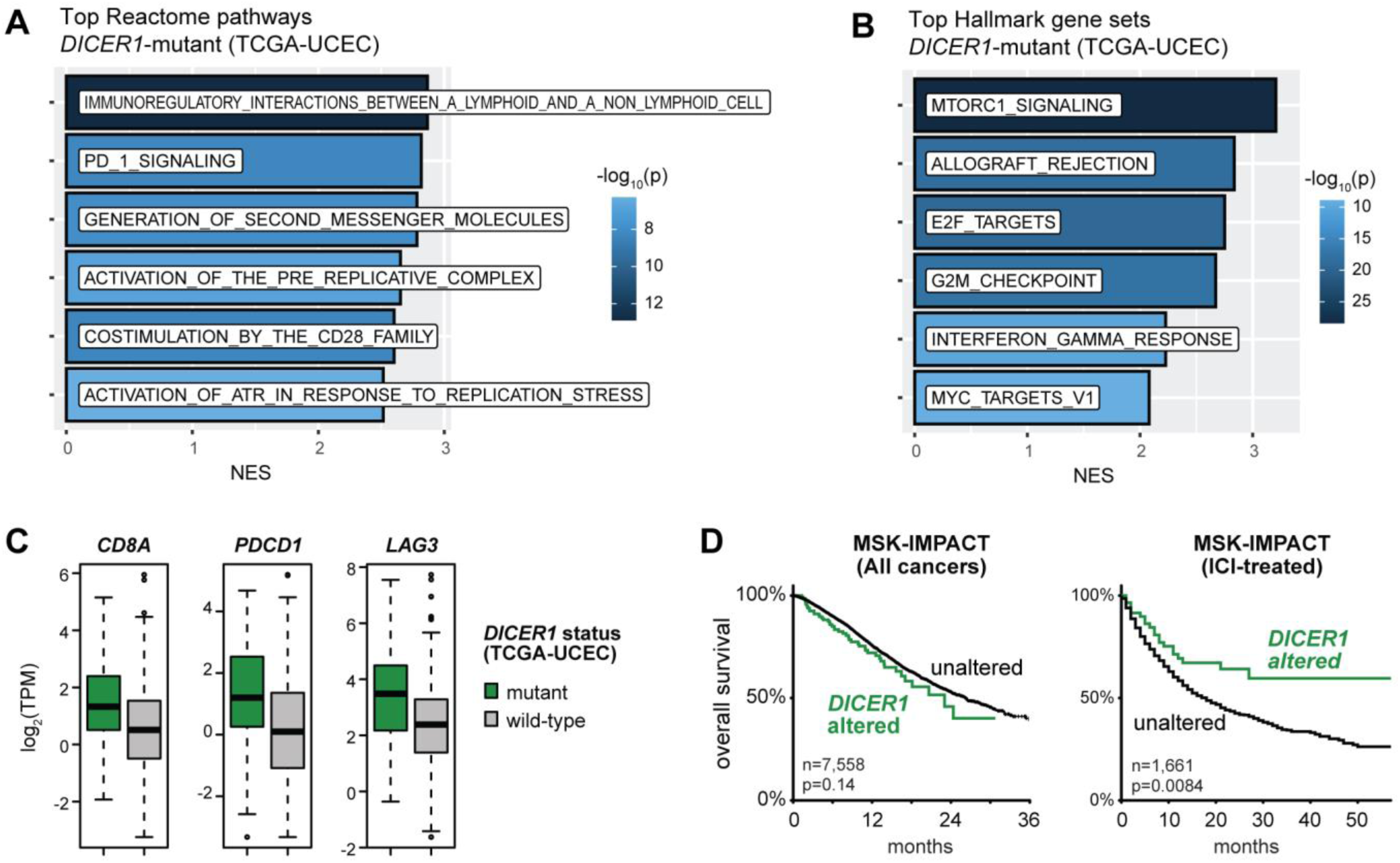
*DICER1* alterations correlate with immune signatures in adult cancer datasets. (A-B) Most enriched Reactome (A) and hallmark (B) gene sets in *DICER1*-mutant vs. *DICER1*-wildtype endometrial cancer. TCGA-UCEC, The Cancer Genome Atlas Uterine Corpus Endometrial Carcinoma. (C) Expression of *CD8A*, *PDCD1*, and *LAG3* in *DICER1*-mutant vs. *DICER1*-wildtype endometrial cancer. TPM, transcripts per million. (D) Clinical outcomes of *DICER1*-altered cancers in MSK-IMPACT regardless of therapy or treated with ICI.

Next, we examined lung adenocarcinoma, the most common type of non-small cell lung cancer. While *DICER1* is rarely mutated in these cancers, lower *Dicer1* expression is associated with tumor progression in mouse models of lung cancer^62^. We thus examined tumors in the bottom 5% by *DICER1* expression, and we found that these were significantly enriched for the same hallmark immune gene sets (**Suppl. Fig. S8B**). As these tumor samples are a mix of tumor and stromal cells, we also asked whether the same patterns arise in cancer cell lines. We queried publicly available RNA-seq results from the 83 lung adenocarcinoma cell lines^63^. Within these 83 lines, we again found that cell lines with lowest *DICER1* expression were enriched for the same immune gene sets (**Suppl. Fig. S8B**). Thus, *DICER1* impairment is associated with an immune-activated expression signature in other tumor types.

Lastly, since the tumor-immune interactions upregulated in *DICER1*-mutant cancers are blocked by clinically available ICIs, we investigated whether mutations in *DICER1* are associated with ICI treatment response. In the pan-cancer MSK-IMPACT cohort^10^, *DICER1* alterations were not linked to improved survival (**Fig. 5C**). In contrast, in the 1,661 patients treated with ICI therapy^64^, *DICER1* mutations were associated with significantly improved survival (**Fig. 5D**). (*DROSHA* was not uniformly profiled in the MSK-IMPACT cohort.)

## DISCUSSION

Single-agent immune checkpoint inhibition has shown minimal efficacy in unselected pediatric solid tumors^4–6^, but it is unknown whether certain subsets of disease may be more likely to respond. Through an unbiased approach, we found several clinical and genomic features of Wilms tumor that were associated with higher levels of PD-L1 and other immune markers. Specifically, we found that chemotherapy treatment is associated with higher levels of PD-L1 in both human tumors and cell lines. Furthermore, mutations in microRNA processing or *CTNNB1* were associated with immune signatures in Wilms tumor specimens. Manipulating these pathways in Wilms tumor cell lines led to PD-L1 upregulation *in vitro*. Lastly, *DICER1* mutations were associated with immune gene signatures and ICI response in adult cancers.

Protein expression analysis revealed that Wilms tumor samples fell into two large groups: one distinguished by immune activation markers, and another by cell cycle markers. Several other groups have shown a dichotomy between proliferation and antitumor immunity in cancer. Through transcriptomic analysis, Su et al.^65^ recently found that Wilms tumors fell into similar clusters, which they termed immune “infiltrated-like Wilms tumor” (iWT) and “desert-like Wilms tumor” (dWT). By gene set enrichment, iWT was enriched for immune-related gene sets, while dWT was enriched for gene sets associated with proliferation, including “DNA repair” and “chromatin-modifying enzymes”. Furthermore, they showed that inhibitors of chromatin-modifying enzymes (specifically, histone deacetylases and EZH2) could enhance the induction of PD-L1 by IFN-γ in 17.94 cells. Lastly, inhibiting cyclin-dependent kinases 4 and 6, which regulate progression through the G1/S cell cycle checkpoint, has also been shown to enhance antitumor immunity *in vivo*^66–68^.

Prior studies showed that PD-L1 staining correlates with worse histology and worse outcomes in Wilms tumor^31,32,69–73^. Our study was designed to discover novel correlations between clinical/genomic features and proteomic signatures, and it lacks the sample size to confirm a significant correlation between such signatures and outcome. However, these prior studies lacked insight into the driving forces behind PD-L1 upregulation in Wilms tumor. Our study identifies a new genomic-phenotypic correlation of PD-L1 elevation in Wilms tumors with *CTNNB1* mutations or microRNA processing mutations. Certain cancers with low mutational burden can still respond to checkpoint blockade if they exhibit mutations that drive PD-L1 overexpression. For instance, tumors can still respond to ICIs despite low TMB if they exhibit inactivation of individual mutations in genes that regulate chromatin remodeling (*ARID1A*, *PBRM1*, *SMARCB1*), RNA processing (*ADAR1*), or immune response (*B2M*, *JAK1/2*)^5,7,8,19–28^. These genes may be important in silencing certain genes that promote immune recognition, and our findings suggest that certain mutational subsets of Wilms tumor may be amenable to immune modulatory therapies despite a low tumor mutational burden.

Our study also shows that chemotherapy can drive PD-L1 expression in Wilms tumor, and this effect may be independent of genomic subtype. This phenomenon has been described in adult cancers, as chemotherapy can induce PD-L1 expression *in vitro*, and the combination of conventional chemotherapy with checkpoint inhibition can produce an added response in clinical trials^7,8,41,74–76^. In other childhood cancers, higher levels of PD-L1 have been noted in rhabdomyosarcomas or osteosarcomas after treatment with chemotherapy^77,78^. Previous studies in Wilms tumors demonstrated immune infiltration after chemotherapy but had not connected chemotherapy with PD-L1 specifically^39,70,73^. Interestingly, the increase in PD-L1 expression we observed varied by chemotherapeutic agent and was not dependent on cell death. The most consistent responses we observed were from doxorubicin, SN-38, and vincristine. Topoisomerase inhibitors, including both anthracyclines like doxorubicin and camptothecins like irinotecan or SN-38, have been previously shown to upregulate PD-L1 in other cancer types^78,79^. In these instances, it is thought that tumor cells induce PD-L1 to evade immunogenic cell death when they experience DNA damage. However, other DNA damaging agents we investigated did not induce similar levels of PD-L1, so other mechanisms may also be involved. Interestingly, we found that vincristine, a chemotherapeutic that acts through microtubule destabilization rather than direct DNA damage, also upregulates PD-L1. Similar findings have also previously been seen in other cancers^42,80^. Regardless, our study adds to the growing literature that make tumor-immune signaling an attractive target for therapy in some pediatric cancer patients, potentially in combination with conventional chemotherapy drugs that are already commonly used.

## METHODS

### Reverse phase protein array

Fifty-three flash-frozen Wilms tumor tissue samples from the Children’s Medical Center biorepository with adequate tissue were sent to the MD Anderson Reverse Phase Protein Array Core for analysis using their standard protocol^30^. Normalized, log2-transformed, median centered values were used for unsupervised hierarchical clustering and plotting. Genomic and transcriptomic analyses for these tumors was previously described^3^. DNA sequencing from tumor and normal samples were aligned to GRCh38 and processed for copy number analysis using cnvKit^81^ (v0.9.5).

### RNA-seq

We used data available through Genomic Data Commons (GDC) release 13.0 (September 2018) to access NCI TARGET Wilms tumor dataset. Reported mutations, copy number data, and RNA-seq expression from TARGET tumors were downloaded from the TARGET data matrix (https://ocg.cancer.gov/programs/target/data-matrix). At any given gene, tumors were designated as having copy number loss or gain when log_2_ copy number was < -0.3 or > +0.3, respectively. These copy number changes were used for mutational classification: copy-number gain of *MYCN* (MYCN); copy-number loss of *REST* (chromatin remodeling); and copy-number loss of *WT1*, *AMER1*, or *RERE* (kidney development). Gene expression quantifications were obtained through the GDC data portal (https://portal.gdc.cancer.gov/). Differential gene expression analysis was performed using DESeq2 (v1.36.0)^82^. The Wald statistic from DESeq2 output then underwent gene set enrichment analysis using fgsea^83^ (v1.10.1) with gene set annotations from MSigDB^84^ (v7.2). To estimate total immune infiltration, we used ESTIMATE default gene signatures^37^. For immune deconvolution, we used QuanTIseq using default parameters^38^.

For adult cancer expression signatures, RNA-seq counts data were downloaded from TCGA (https://www.cancer.gov/tcga) on Dec. 7, 2020. Tumors were categorized as “low *DICER1*” if they were in the bottom 5% of tumors by *DICER1* expression. As above, differential expression analysis was performed using DESeq2 and fgsea. Outcomes for MSK-IMPACT patients were generated from cBioPortal^64,85,86^.

### Tissue culture

All cell lines used were cultured in an incubator at 37°C with 5% CO_2_. WiT49 (RRID: CVCL_0583) was a gift from Sharon Plon’s laboratory; 17.94 was purchased from Ximbio (cat. no. 153333); and JR-1, RD, Kelly, and SHEP were gifts from Stephen Skapek’s laboratory. WiT49, 17.94, and RD were maintained in Dulbecco’s Modified Eagle Medium (DMEM) supplemented with 10% fetal bovine serum (FBS) and antibiotic-antimycotic (Gibco 15240062). JR-1 was maintained in DMEM supplemented with 20% FBS and antibiotic-antimycotic. KELLY and SHEP were maintained in RPMI 1640 supplemented with 10% FBS and antibiotic-antimycotic. Testing for mycoplasma (Lonza LT07-318) was done every 6 months (last negative test for all six lines was February 2024. Cell identity was verified annually by short tandem repeat genotyping (last verified WiT49, JR-1, RD, Kelly, and SHEP in February 2024; 17.94 in April 2024. Negative non-targeting (NTC) controls and knockdowns of *DROSHA* and *DICER1* in WIT49 cells was performed with CRISPR interference (CRISPRi) was previously described^3^. Transduced cells were continuously maintained in 0.5 µg/ml puromycin, with all downstream applications performed at no more than 9 passages.

### Gene expression quantification

Total RNA was extracted from subconfluent cells at low passage using miRNeasy Mini Kit with DNAse I digestion (Qiagen 217400 and 79254). RNA was reverse transcribed with iScript Reverse Transcription Supermix (Bio-Rad 1708841), and we quantified primary and mature transcripts for *CD274* with quantitative PCR (qPCR) with iTaq™ Universal SYBR® Green Supermix (Bio-Rad 1725125). The mature transcript qPCR primers (Forward: TGCAGGGCATTCCAGAAAGA; Reverse: ATAGGTCCTTGGGAACCGTG) span two exons to disfavor unspliced transcripts. Conversely, primers for the primary *CD274* transcript (Forward: TGAAGCAGTCTTCTTTTCGTGT; Reverse: TTACCGTTCAGCAAATGCCA) amplify a region near the 3’ end of an intron and near the 5’ end of the adjacent downstream exon to exclude processed mRNA. For Alu RNA, primers used were described previously^56^ (Forward: CAACATAGTGAAACCCCGTCTCT; Reverse: GCCTCAGCCTCCCGAGTAG). For normalization, we used 18s rRNA (GTAACCCGTTGAACCCCATT, CCATCCAATCGGTAGTAGCG), calculated relative expression using the 2^-ΔΔCt^ method, and determined significance by unpaired two-tailed Student’s T-test versus both NTC cell lines.)

Western blot was done as previously described^3^, using subconfluent WiT49 or 17.94 cells cultured in complete media without puromycin for at least 2 days. For CHIR-99021 treatment, cells were treated with 10µM CHIR 99021 (STEMCELL Technologies, NC1267203) for 3, 6, 24 hours, or equivalent DMSO for 24 hours. Primary antibodies used are as follows: DICER1 (1:3000, Cell Signaling Technology, cat. 5362, RRID:AB_10692484), DROSHA (1:3000, Cell Signaling Technology, cat. 3364, RRID:AB_2070685); PD-L1 (1:3000, Cell Signaling Technology, cat. 13684, AB_2687655); β-catenin (1:1000, Cell Signaling Technology, cat. 8480, RRID:AB_11127855); active β-catenin (1:1000, Cell Signaling Technology, cat. 8814, RRID: AB_11127203); and tubulin (1:3000, Cell Signaling Technology, cat. 3873, RRID:AB_1904178). Each run was performed at least twice to ensure reproducibility.

### Flow Cytometry

After cells were seeded, media was replaced with drugs in fresh media at concentrations indicated. These drugs were as follows: doxorubicin (Fisher Scientific, D419325MG), vincristine (Thermo Fisher, J60907.MA), cyclophosphamide (R&D Systems, 4091-50), SN-38 (MedChem Express, HY-13704), temozolomide (MedChem Express, HY-17364), actinomycin D (Sigma Aldrich, A9415), and CHIR 99021 (STEMCELL Technologies, NC1267203). One well each received recombinant human IFNγ (Pepro Tech, 300-02) as a positive control; DMSO alone as a vehicle control; and media alone as a viability control. Cells were treated for 24 hours before being collected for flow cytometry with TrypLE (Gibco, 12604-013). For some experiments, cells were fixed prior to flow cytometry by resuspending in 4% formaldehyde at room temperature for 15 minutes. As a viability control, cells were heat-killed in a 65°C water bath for 20 mins. Cells were stained with PE-conjugated anti-PD-L1 antibody (Thermo Fisher, cat. no. 12-5983-42) at a dilution of 1:100 for 30 min. on ice. Afterwards, cells were washed and resuspended in buffer with the viability stain 7-aminoactinomycin D (7-AAD, Thermo Fisher, cat. no. 00-6993-50) before analysis by flow cytometer (NovoCyte Advanteon). Data was analyzed using FlowJo software (version 10). Each run was performed at least twice to ensure reproducibility.

### Fluorescent Immunocytochemistry

Cells were seeded into 8-chamber slides (Ibidi 80827 or Falcon 354118) in complete culture media without puromcyin. After 24 hours, for a PD-L1 positive control, a chamber with WiT49 + sgNTC-2 cells were replaced with complete media with 20 ng/ml IFNγ. For a dsRNA positive control, WiT49 + sgNTC-2 cells were transfected with 10 µg/ml polyinosinic-polycytidylic acid (Poly I:C) with Lipofectamine 3000 (Invitrogen L3000015). The next day, cells were fixed with 4% formaldehyde; permeabilized with 0.5% Triton-X100; and blocked with 5% donkey serum. The primary antibodies used were PD-L1 (Invitrogen 14-5982-82, diluted 1:50) or anti-dsRNA antibody (Sigma MABE1134, clone rJ2, diluted 1:60). The secondary antibody used was donkey anti-mouse IgG conjugated to Alexa Fluor 488 (Invitrogen A-21202, diluted 1:10,000). After counter staining with DAPI, slides were imaged using the Laser scanning confocal LSM880 with Airyscan (Zeiss) and ZEN Microscopy Software (Zeiss).

## ACKNOWLEDGMENTS

We thank the patients and families who contributed to this study. This work was financially supported by grants from the Pablove Foundation (Childhood Cancer Research Seed Grant to P.D.B.T.), Alex’s Lemonade Stand Foundation (Young Investigator grant to K.S.C.), Cancer Prevention and Research Institute of Texas (RR180071 to K.S.C.), the National Cancer Institute (K08CA207849, P50CA196516, P30CA142543, and R01CA289259 to K.S.C.), and the U.S. Department of Defense (KC220019 to K.S.C.). This research was also supported by the computational resources provided by the BioHPC supercomputing facility in the Lyda Hill Department of Bioinformatics at UT Southwestern. The Functional Proteomics Reverse Phase Protein Array Core is supported in part by The University of Texas MD Anderson Cancer Center and the National Institutes of Health (P30CA016672 and R50CA221675). The authors would like to acknowledge the Quantitative Light Microscopy Core, a Shared Resource of the Harold C. Simmons Cancer Center, supported in part by an NCI Cancer Center Support Grant, 1P30CA142543, and specifically, use of the Laser scanning confocal Zeiss LSM880 inverted with Airyscan (1S10OD021684-01 to Kate Luby-Phelps). We would also like to thank Andrew Koh for sharing access to laboratory equipment, Stephen Skapek for sharing cell lines, and James Amatruda for mentorship and support. The results shown here are in part based upon data generated by the TCGA Research Network (https://www.cancer.gov/tcga) and the Therapeutically Applicable Research to Generate Effective Treatments (https://www.cancer.gov/ccg/research/genome-sequencing/target) initiative, phs000218. The data used for this analysis are available at the Genomic Data Commons (https://portal.gdc.cancer.gov).

## SUPPLEMENTARY FIGURE LEGENDS

**Supplementary Figure S1.**
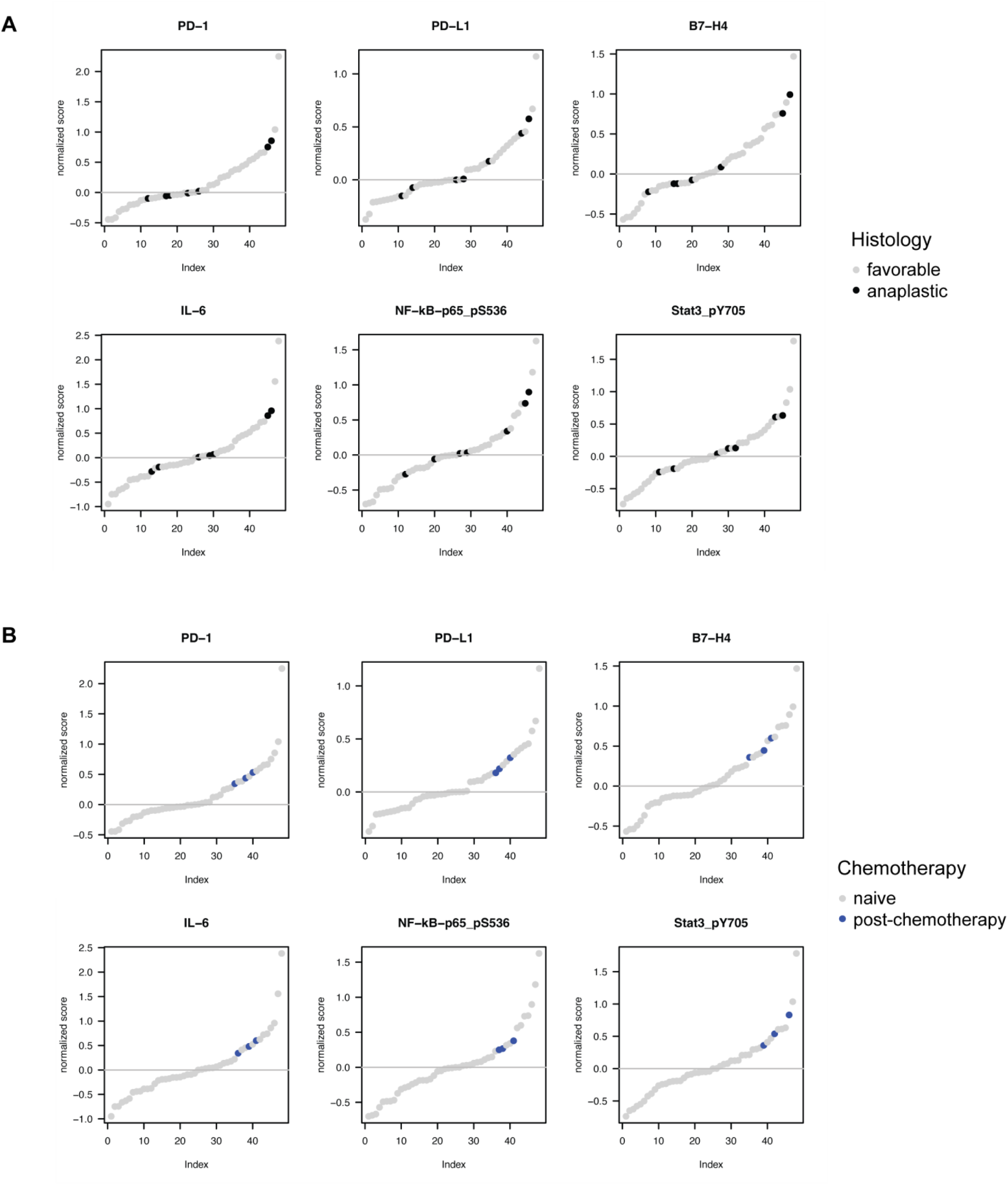
Correlation of clinical features with RPPA expression of individual immune markers. (A-B) RPPA expression of PD-1, PD-L1, B7-H4, IL-6, phospho-NF-κB, and phospho-Stat3, based on histology (A) and chemotherapy status (B).

**Supplementary Figure S2.**
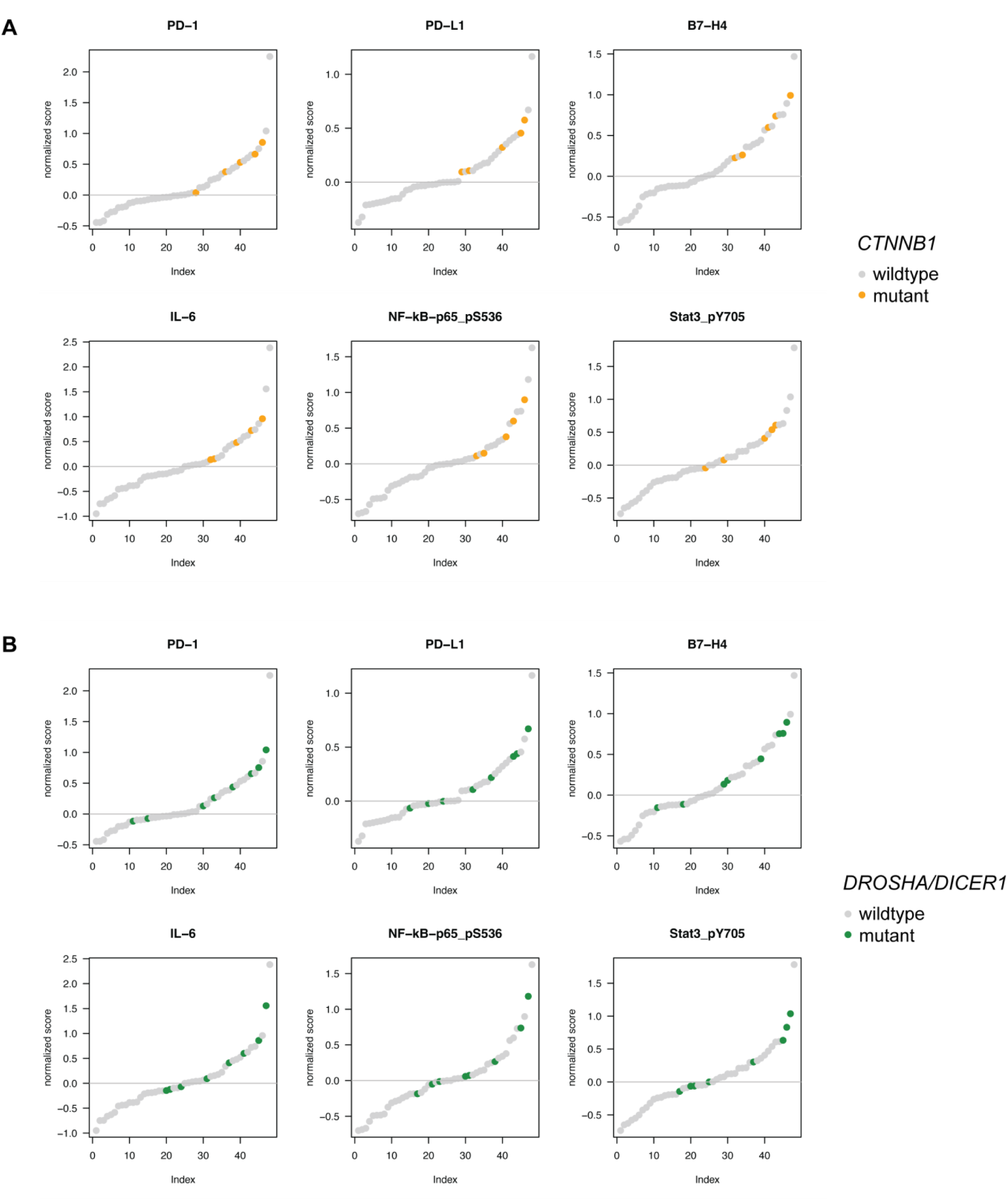
Correlation of genomic features with RPPA expression of individual immune markers. (A-B) RPPA expression of PD-1, PD-L1, B7-H4, IL-6, phospho-NF-κB, and phospho-Stat3, based on mutation in *CTNNB1* (A) and *DROSHA/DICER1* (B).

**Supplementary Figure S3.**
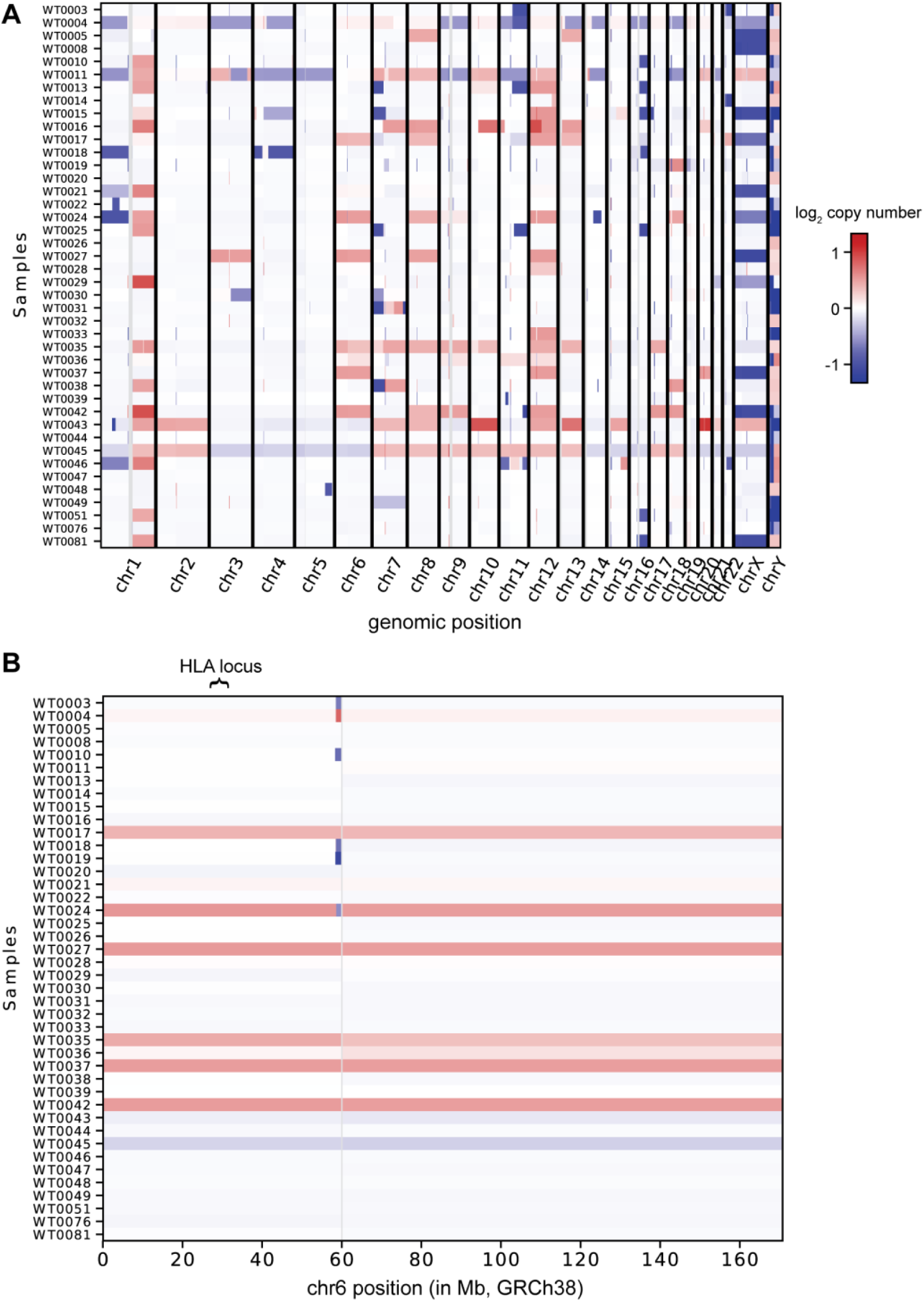
Copy number changes in profiled Wilms tumors. (A) Genome wide copy number changes in Wilms tumors profiled in this study. (B) Copy number changes in chr6 in Wilms tumors profiled in this study.

**Supplementary Figure S4.**
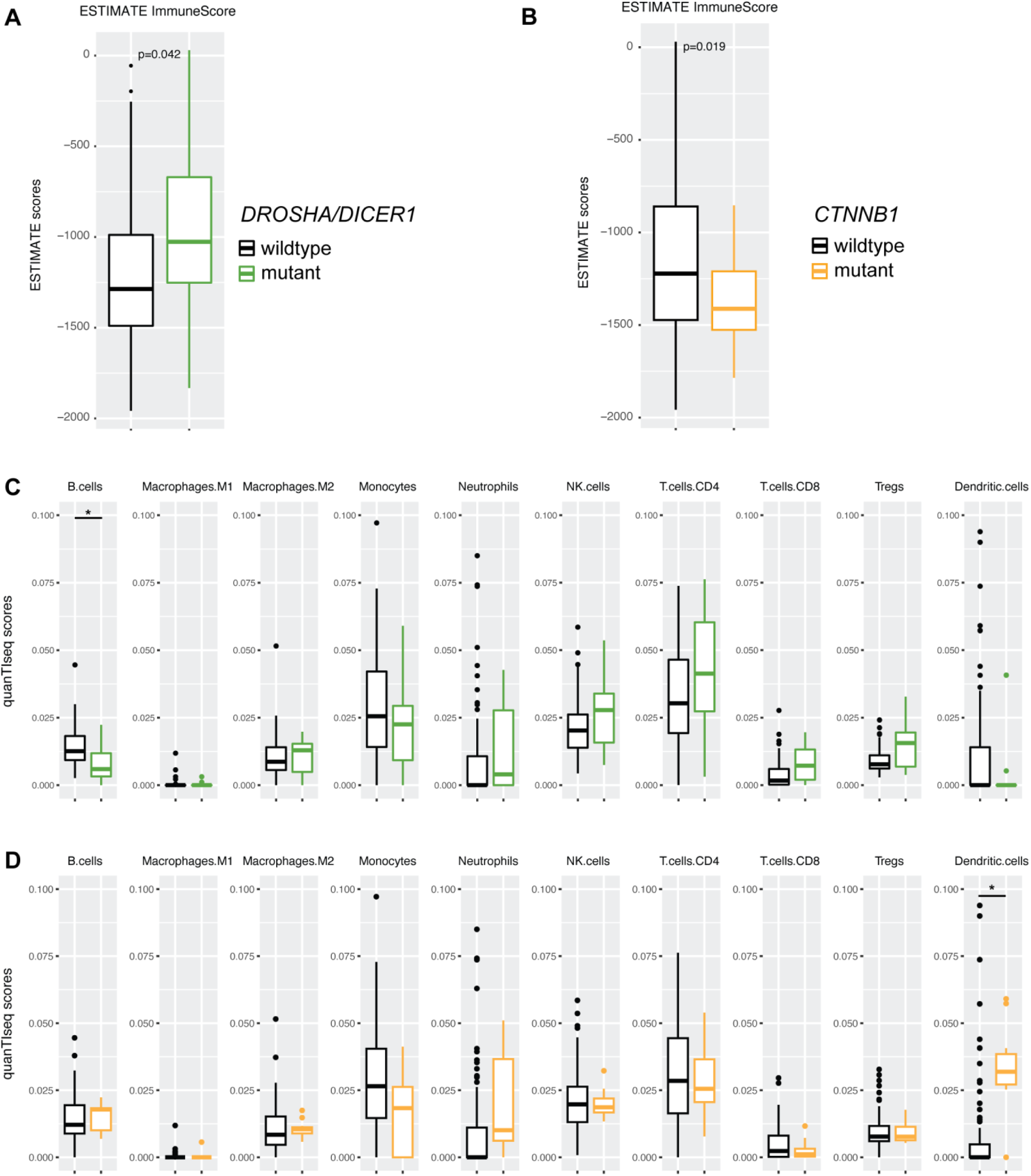
*DROSHA*-mutant Wilms tumors in TARGET dataset exhibit a more inflammatory transcriptome. (A-B) ESTIMATE immune score, classified by mutation in microRNA processing genes (A) or *CTNNB1* (B). (C-D) Immune cell proportions based on quanTIseq, classified by mutation in microRNA processing genes (C) or *CTNNB1* (D). (*adjusted p value < 0.05, computed by Student’s t test and adjusted by Benjamini-Hochberg method for multiple comparisons)

**Supplementary Figure S5.**
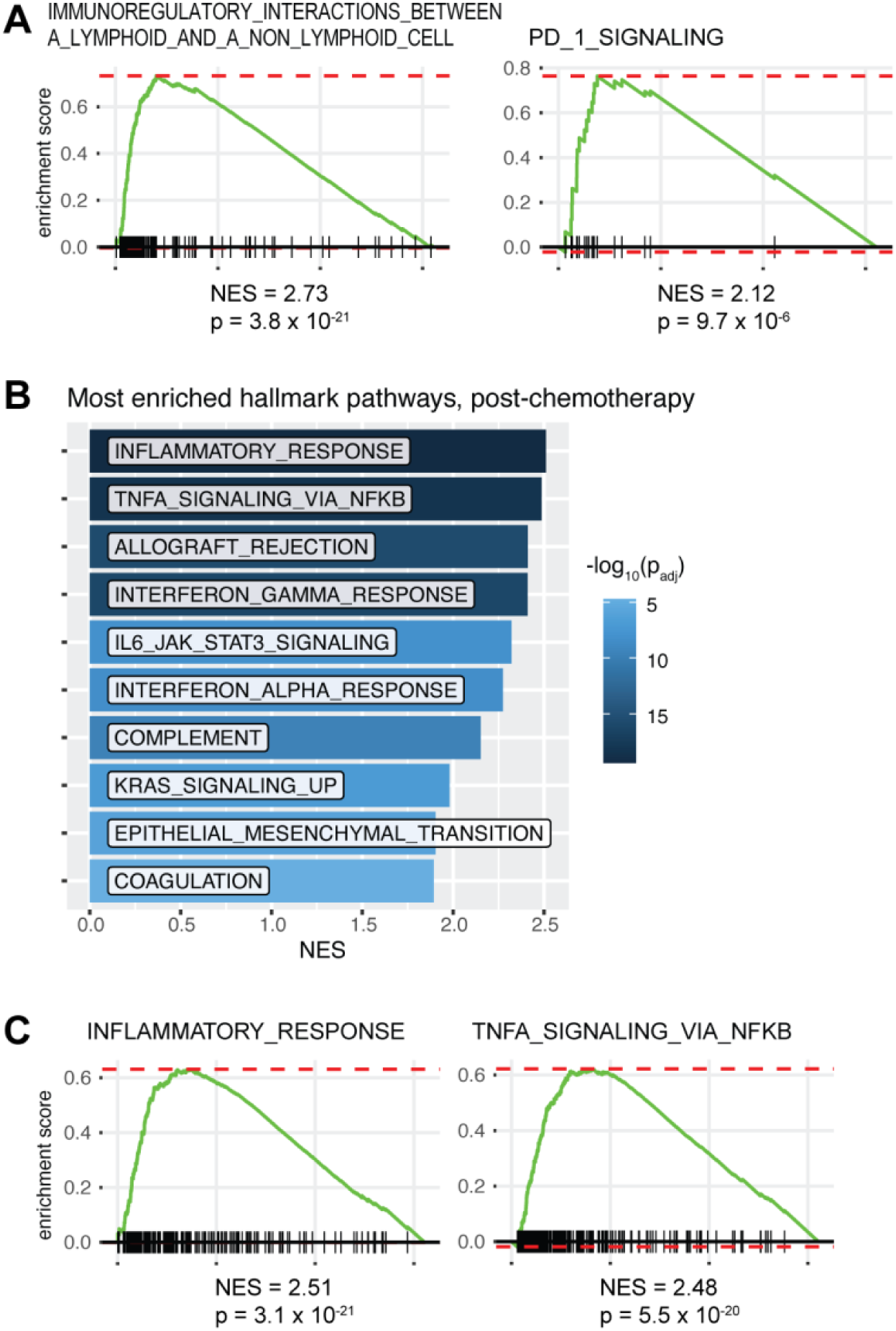
Chemotherapy induces immune signatures in Wilms tumor RNA-seq. (A) GSEA enrichment for top two immune-related Reactome gene sets in chemotherapy-treated tumors (from US) vs. chemotherapy-naïve tumors (from Europe). (B) Most enriched hallmark gene sets in chemotherapy-treated tumors. (C) GSEA enrichment for top two immune-related hallmark gene sets in chemotherapy-treated tumors.

**Supplementary Figure S6.**
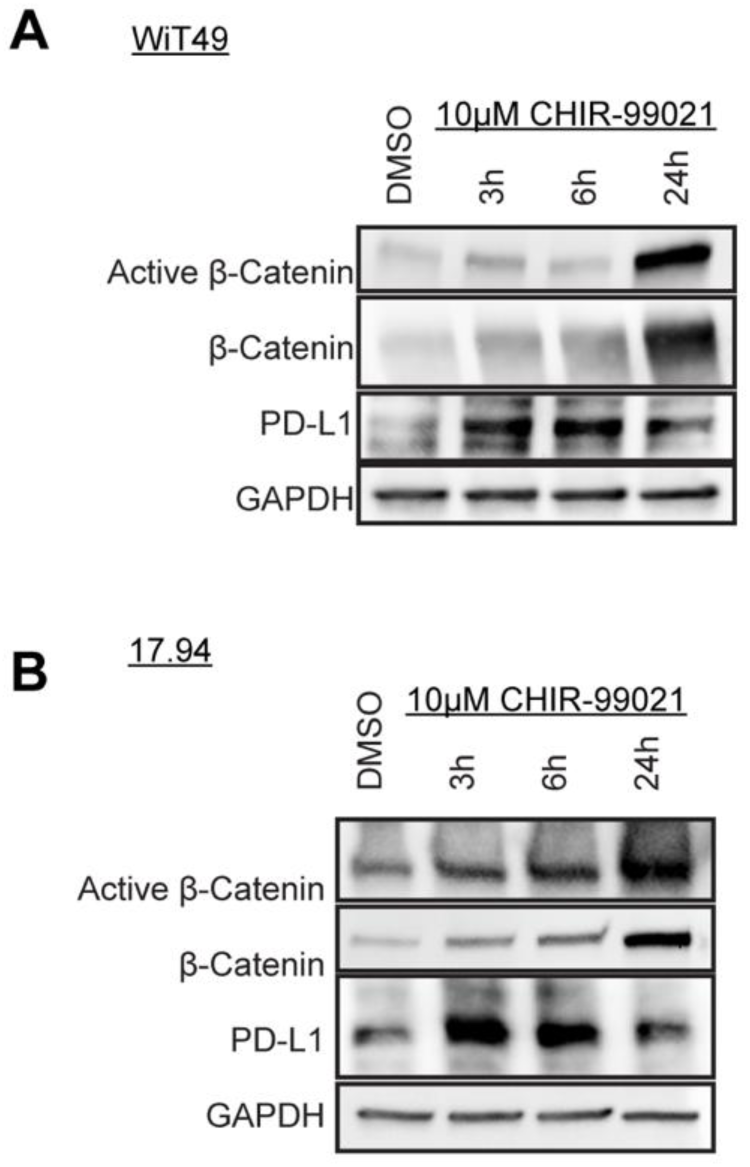
CHIR-99021 upregulates PD-L1 in Wilms tumor cells. (A-B) Western blot of WiT49 (A) and 17.94 (B) treated with vehicle or 10µM CHIR-99021 for 3, 6, and 24 hours.

**Supplementary Figure S7.**
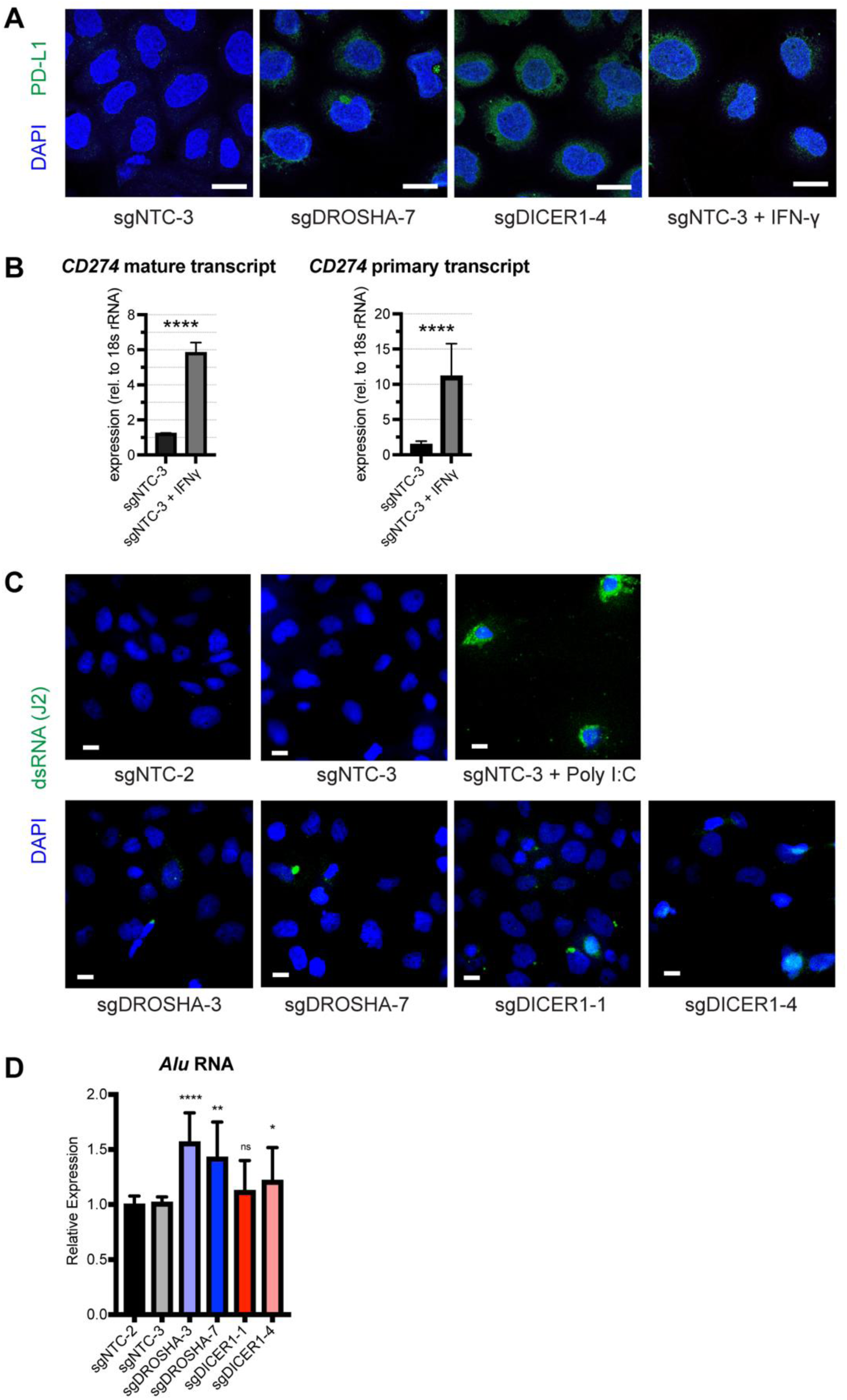
*DROSHA* and *DICER1* knockdowns increase PD-L1. (A) Representative images of PD-L1 immunocytochemistry of WiT49 *DROSHA* and *DICER1* knockdowns versus NTC (Scale bar = 25µm). (B) Quantitative PCR of *CD274* primary and mature transcripts in WiT49 NTC cells following interferon-γ (IFNγ) treatment. (****p<0.0001 by unpaired two-tailed Student’s t-test) (C) Fluorescent immunocytochemistry of dsRNA in WiT49 cells with *DROSHA/DICER1* knockdown. Scale bar = 50µm. Transfection with polyinosinic:polycytidylic acid (poly I:C) used as positive control. (D) Quantitative PCR of *Alu* RNA in WIT49 cells. (****p<0.0001, **p<0.01, *p<0.05, ns p ≥ 0.05, by unpaired two-tailed Student’s t-test versus NTC cell lines)

**Supplementary Figure S8.**
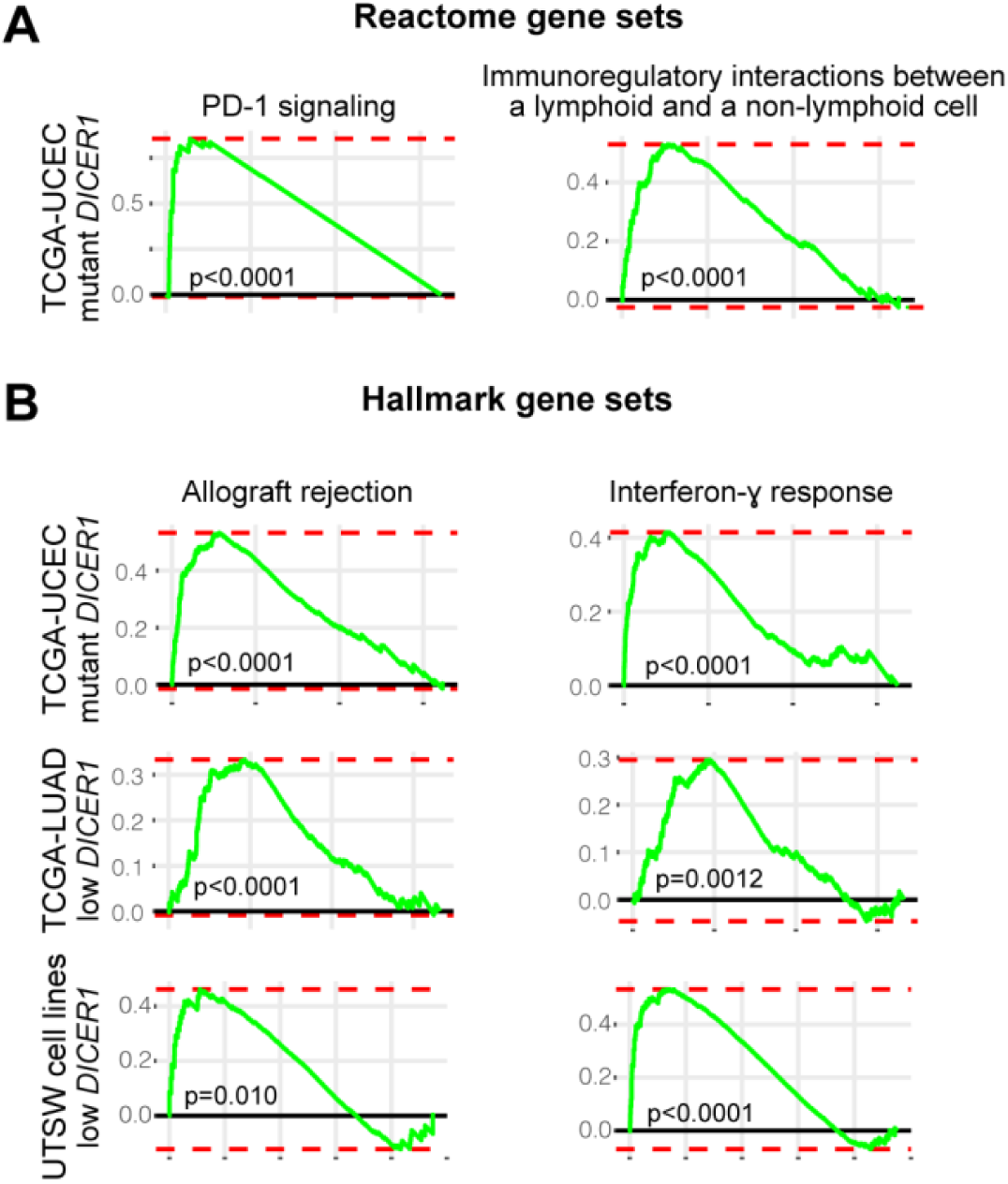
Immune signatures enriched in adult cancer datasets with *DICER1* impairment. (A) GSEA plots of immune-related Reactome gene sets in *DICER1*-mutant endometrial cancers highlights T cell-cancer interactions. (B) GSEA plots of allograft rejection and interferon gamma response gene sets in TCGA datasets. From top to bottom: *DICER1*-mutant endometrial cancer (UCEC); the bottom 5% of lung adenocarcinoma (LUAD) by *DICER1* expression; and the bottom 5% of UTSW lung adenocarcinoma cell lines by *DICER1* expression.

## SUPPLEMENTARY TABLES

**Supplementary Table S1.** Clinical and molecular features from the Wilms tumor patients profiled in this study.

**Supplementary Table S2.** Normalized RPPA expression from the Wilms tumors profiled here.

